# Dopamine in the basal amygdala signals salient somatosensory events during fear learning

**DOI:** 10.1101/716589

**Authors:** Wei Tang, Olexiy Kochubey, Michael Kintscher, Ralf Schneggenburger

## Abstract

The amygdala is a brain area critical for the formation of threat memories. However, the nature of the teaching signal(s) that drive plasticity in the amygdala are still under debate. Here, we use optogenetic methods to investigate whether dopamine release in the amygdala contributes to fear learning. Antero- and retrograde labeling showed that a sparse, and relatively evenly distributed population of ventral tegmental area (VTA) neurons projects to the basal amygdala (BA). *In-vivo* optrode recordings in behaving mice showed that many VTA neurons, amongst them putative dopamine neurons, are excited by footshocks. Correspondingly, *in-vivo* fiber photometry of dopamine in the BA revealed robust dopamine concentration transients upon footshock presentation. Finally, silencing VTA dopamine neurons, or their axon terminals in the BA during the footshock, reduced the extent of threat memory retrieval one day later. Thus, VTA dopamine neurons projecting to the BA code for the saliency of the footshock event, and the resulting dopamine release in the BA facilitates threat memory formation.

## Introduction

Animals must predict dangers in their environment to ensure survival; for this reason, powerful mechanisms of harm detection and fear learning have evolved in animals (Feinberg and Mallatt, 2017). During auditory cued fear learning, an innocuous sensory percept like a tone (the conditioned stimulus, CS), acquires a negative emotional valence, and will elicit a defensive behavior after pairing with an aversive outcome, like a mild footshock (the unconditioned stimulus, US) (LeDoux, 2000; Tovote et al., 2015). The amygdala has been identified as a brain structure with an important role in fear learning (Davis, 1992; LeDoux, 2000; Duvarci and Paré, 2014; Tovote et al., 2015). Action potential (AP) firing in response to a tone is increased in many lateral amygdala (LA) and BA neurons during - and after fear conditioning protocols (Quirk et al., 1995; Amano et al., 2011; Grewe et al., 2017). It is thought that synapses in the LA that code for the CS undergo LTP, and then drive increased LA neuron firing upon CS presentation (Rumpel et al., 2005; Sigurdsson et al., 2007; Nabavi et al., 2014). The nature of the synaptic- and/or neuromodulatory signal that instructs plasticity in the LA and BA is currently investigated (Herry and Johansen, 2014). This “teaching signal” for plasticity is likely represented, on the one hand, by depolarizing inputs coding for the US, which can induce associative plasticity. Indeed, footshock stimulation causes cfos expression in a sparse, but distinct neuronal population in the LA and BA (Gore et al., 2015). Furthermore, we have recently identified a glutamatergic synaptic input from the insular cortex as a candidate pathway for one such “depolarizing” teaching signal in the LA (Berret et al., 2019).

In addition to direct glutamatergic depolarization of LA- and BA neurons, behaviorally salient stimuli like footshocks recruit several neuromodulatory systems, which, amongst other target areas, project to the BA and LA where they can facilitate the acquisition of threat memories (Johansen et al., 2014). Recent work has shown that both acetylcholine and noradrenaline, acting in the LA and/or BA, facilitate the formation of fear memories (Jiang et al., 2016; Uematsu et al., 2017). We wished to investigate whether dopamine acting in the amygdala might be an additional neuromodulator that facilitates the formation of threat memories.

Dopamine neurons in the midbrain ventral tegmental area (VTA) are a heterogenous population that project to the prefrontal cortex, the Nucleus acumbens (NAc) and the amygdala (Asan, 1998; Wise, 2004); (Lammel et al., 2011; Lammel et al., 2012; Beier et al., 2015), and the different dopaminergic projections can be involved in both appetitively and aversively motivated behavior. Early evidence for a role of dopamine signaling in the amygdala during aversive learning came from measurements of enhanced dopamine in the basolateral amygdala and in the NAc after stressful stimuli (Young and Rees, 1998; Inglis and Moghaddam, 1999; de Oliveira et al., 2011), and from infusion of dopamine receptor antagonists in the amygdala that reduced fear learning (Lamont and Kokkinidis, 1998; Guarraci et al., 1999; Nader and LeDoux, 1999; Heath et al., 2015). However, classical studies in monkeys and rats have found a role for VTA dopamine neurons in reward processing (Schultz, 1998; Wise, 2004), and some studies found that VTA dopamine neurons are inhibited by aversive stimuli (Mirenowicz and Schultz, 1996; Schultz, 1998; Ungless et al., 2004; Tan et al., 2012). Thus, a role of dopamine neurons in aversively motivated behavior has remained somewhat controversial, despite evidence from genetic manipulations indicating a role of the dopamine system in aversive learning (Fadok et al., 2009; Zweifel et al., 2011), and despite findings that some dopamine neurons in the VTA are excited by aversive stimuli (Guarraci and Kapp, 1999; Brischoux et al., 2009; Gore et al., 2014); see (Horvitz, 2000), for an insightful early review). In the present study, we have used optogenetic methods, *in-vivo* optrode recordings, *in-vivo* fiber photometry and circuit mapping techniques to investigate the role of a VTA to BA dopaminergic projection in auditory cued fear learning.

## Materials and Methods

### Animals

All procedures with laboratory animals (*mus musculus*) were authorized by the veterinary office of the Canton of Vaud, Switzerland (authorizations 2885.0 and 3274.0). The following mouse lines were used in this study: 1) DAT-internal ribosome entry site (IRES)-Cre line (B6.SJL-Slc6a3^tm1.1(cre)Bkmn^/J, Jackson Lab #006660) called here *DAT*^*Cre*^ mice (Bäckman et al., 2006); 2) Cre-dependent channelrhodopsin 2 reporter line (Rosa26:lsl:ChR2-eYFP, Jackson Lab #024109, also known as Ai32) called here *ChR2-eYFP* mice (Madisen et al., 2012); 3) C57Bl6/J wild type mice (Jackson Lab #000664). All experimental mice were virgin males, group-housed under a 12/12 hr light/dark cycle with food and water ad libitum until separated into single cages one day before surgery.

### Viral vectors

For Cre-dependent expression of Arch in DAT^+^ neurons (Figs. 3, 4), an AAV1:CBA:FLEX:Arch-GFP vector was used (UPenn [Addgene], cat. no. AV-1-PV2432 [22222-AAV1]). In mice for the control groups (Figs. 3, 4) and for anterograde tracing (Fig. 1), eGFP expression was driven in DAT^+^ neurons by an AAV1:CAG:FLEX-eGFP vector (UPenn [Addgene], cat. no. AV-1-ALL854 [51502-AAV1]). For *in-vivo* fiber photometry of dopamine release (Fig. 5), an AAV5:hsyn:dLight1.2 vector was used (Addgene; cat. no. 111068-AAV5).

**Fig. 1.**
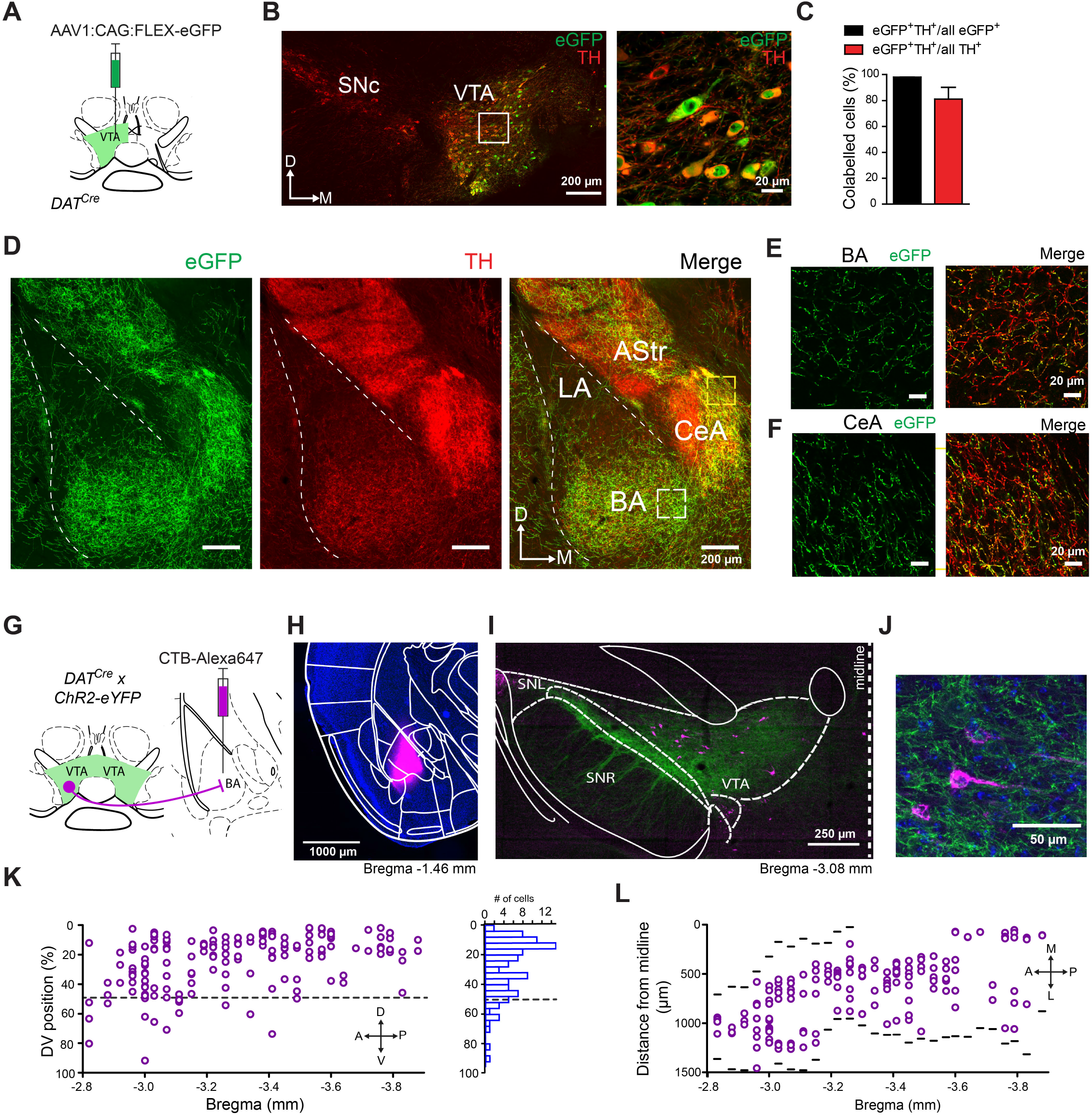
A dopaminergic afferent projection from the VTA to different amygdala subnuclei (**A**) Schematic of the experimental approach for panels B - F. An AAV1 vector driving the Cre-dependent expression of enhanced green fluorescent protein (eGFP) virus was injected into the VTA of a DAT^Cre^ mouse. (**B**) The injection area on the level of the VTA, with eGFP fluorescence (green channel) and the immunohistochemistry with an anti-TH antibody (red channel). The area indicated by the white box is shown at a higher magnification on the right. Note that eGFP expression is limited to TH^+^ cells. (**C**) Quantifications of, *left*, the percentage of eGFP^+^ cells that were also TH^+^ amongst all eGFP^+^ cells, and, *right*, the percentage of eGFP^+^ *and* TH^+^-positive cells within all TH^+^ cells, respectively (mean ± S.E.M; from n = 9 VTA sections from n = 3 mice). (**D**) eGFP-expressing axons were observed in the basal amygdala (BA), medial portion of the central amygdala (“CeA” labels the entire central amygdala) and in the amygdala-striatal transition zone (AStr), but were largely absent in the lateral amygdala (LA). (**E - F**) Confocal images of eGFP and TH^+^ fibers in the BA (E) and CeA (F). (**G**) Schematic of the retrograde labelling approach for panels H-L. A fluorescently labeled cholera toxin B (CTB-Alexa 647, magenta) was stereotaxically injected into the BA of a DAT^Cre^ × ChR2-eYFP mouse. (**H**) Example image of the injection site in the BA, overlaid with the outlines (white lines) of the brain areas from the mouse brain atlas (Franklin and Paxinos, 2016). (**I**) Image of a coronal brain section containing the VTA at the indicated Bregma position. The magenta channel shows neurons back-labelled with CTB-Alexa 647; green channel shows expression of a ChR2-eYFP reporter gene in DAT^+^ neurons. (**J**) A confocal image of a CTB-Alexa 647 labelled BA-projecting neuron in the VTA. (**K**) Dorso-ventral (DV) position of CTB-Alexa 647 - positive neurons within the VTA (0% represents most dorsal position), for all sections along the AP axis of one mouse, and the corresponding histogram (right). Note the preferential position of BA projectors in the dorsal half of the VTA. (**L**) Medio-lateral position of CTB-Alexa 647 - positive neurons, again for all sections along the AP-axis. The black lines show the medial or lateral border of the VTA; from Bregma ~ −3.3 mm the bilateral nuclei of the VTA meet at the midline. Note that there was no apparent preferential localization of BA projectors on the medio-lateral axis, nor on the anterior - posterior axis. Similar observations were made in another mouse.

### Virus injection and optic fiber implantation

The age of the mice at surgery was 42-49 days. For *in-vivo* optogenetic silencing experiments virus was injected into the VTA of *DAT*^*Cre*^ mice, and optic fibers were implanted unilaterally above the VTA (Fig. 3), or else bilaterally above the BA or bilaterally above the central amygdala (CeA) during the same surgery session (Fig. 4). The fiber coordinates were: AP −1.66 mm, ML ±3.35 mm, DV 3.65 mm for BA, or AP −1.36 mm, ML ±2.85 mm, DV 3.65 mm for CeA (DV measured from the dura surface). The animals were randomly assigned to a “control group” or to an “Arch group”, injected respectively with AAV1:CAG:FLEX-eGFP or AAV1: CBA:FLEX:Arch-GFP virus (see above). All other procedures were the same between the control - and the test groups. Stereotaxic surgeries with a model 940 stereotactic instrument (Kopf Instruments, USA) followed the procedures described in Berret et al. (2019). Virus suspension (200 - 400 nl) was injected unilaterally into the VTA at coordinates relative to bregma: AP −3.3 mm; ML −0.35 mm; DV 4.2-4.6 mm (DV measured from the dura surface).

Fiber optic implants for *in-vivo* optogenetic silencing were custom made from a 200 μm core / 0.39 NA / 230 ◻Photometry optic fibers were made from a 400Photometry optic fibers were made from a 400Photometry optic fibers were made from a 400Photometry optic fibers were made from a 400m outer diameter optic fiber (FT200EMT; Thorlabs) and 1.25 mm outer diameter ceramic ferrules (CFLC230; Thorlabs) as described (Sparta et al., 2012). Yellow light was produced by a 561 nm diode pumped solid state (DPSS) laser (MGL-FN-561-AOM-100mW, CNI Lasers, China) equipped with an AOM and an additional mechanical shutter (SHB05T; Thorlabs) as described previously (Berret et al., 2019). The laser power was adjusted to 10 mW after the implant tip for each individual animal.

The optrode implantation procedure into the VTA (Fig. 2) was similar to the VTA fiber implantation, with an additional stainless steel microscrew (Antrin Miniature Specialties, CA, USA) with a soldered copper ground wire implanted into the skull.

**Fig. 2.**
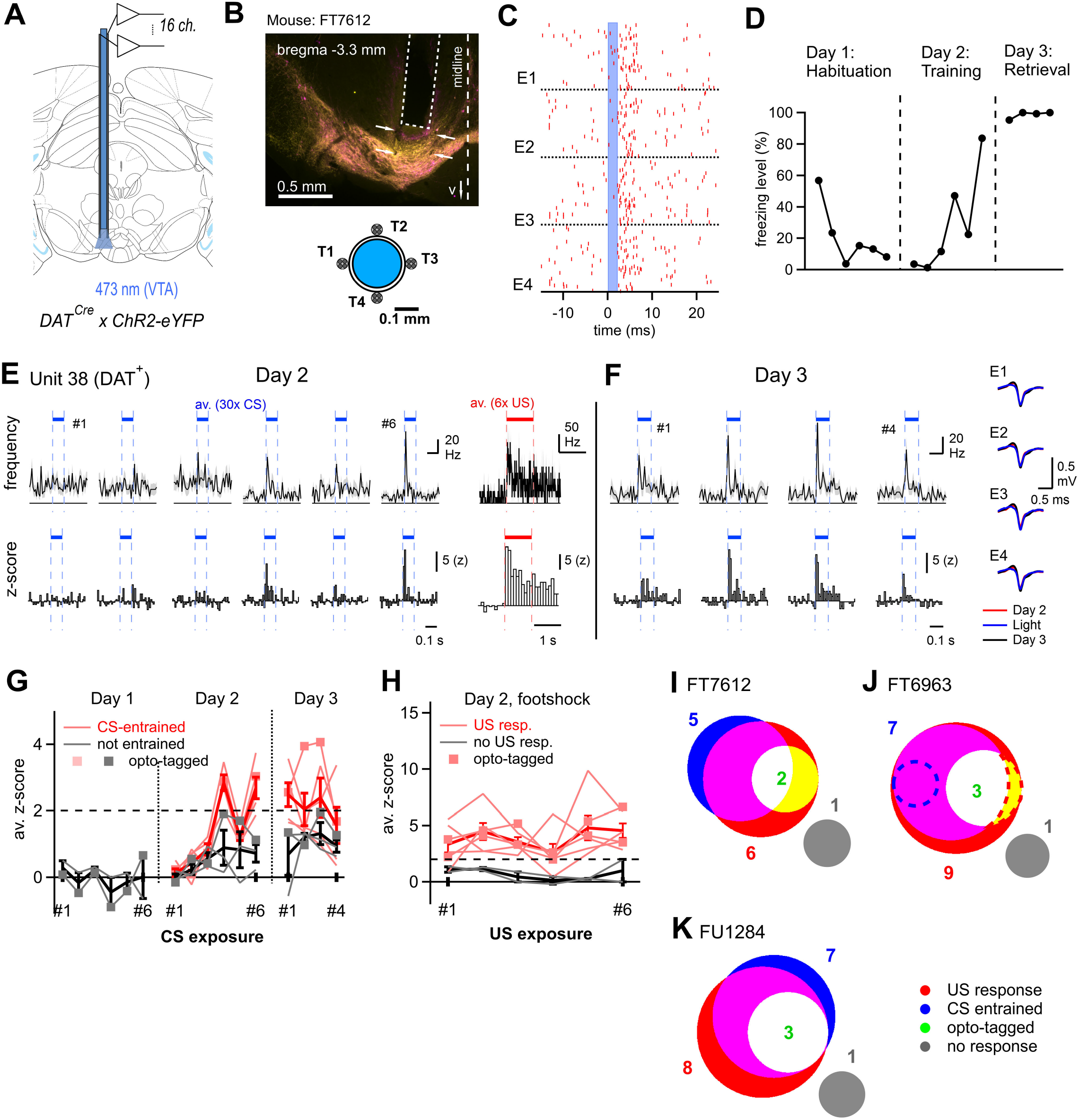
VTA neurons, and amongst them dopamine neurons, respond to footshocks and acquire CS - responsiveness. (**A**) Schematic showing an optrode with 16 recording channels implanted in the VTA of a DAT^Cre^ × ChR2-eYFP mouse. (**B**) Post-hoc histological image showing the placement of an optrode (dashed line) in the VTA in the example mouse of panels B – H (FT7612). The tracks of two tetrodes are visible (arrows). Bottom, scheme of the design of four tetrodes (T1 - T4) around the optical fiber. (**C**) Illustration of optogenetic identification of putative DAT^+^ units. Raster plot for four electrodes (E1-E4) of one tetrode, showing unsorted spikes aligned to the onsets of n = 100, 2 ms – long laser light pulses (blue shading). Spikes at 2-8 ms after the light pulse were collected and subjected to spike clustering (see Materials and Methods). (**D**) Time course of freezing of the example mouse during the three days of the fear learning protocol. (**E**) Spiking activity of a single opto-tagged putative DAT^+^ unit during day 2 (training day). AP frequency (top) and z-score (bottom) are shown in response to the CS (averages over the n = 30 tone presentations for each tone block), as well as in response to footshocks (averages over the n = 6 footshock presentations; right). (**F**) Spiking activity of the same unit as (E), but for responses to CS presentations on day 3 (threat memory retrieval). Right, AP-waveforms for the unit shown in E, F. (**G, H**) Responses of all units in this mouse (FT7612) to tone presentations during day1 - day 3 (G), and to footshocks on day 2 (H). Z-scores were analyzed and plotted as a function of time (number of tone block). Units classified as CS - entrained are shown in pink, and the others in gray. Thick red and black lines show average ± S.E.M. across these groups, respectively. Square symbols connected by lines represent DAT^+^ units identified by optotagging. (**I, J, K**) Venn-type diagrams showing the number of units in the different response classes and their overlap. The data in (I) is from the example mouse shown in panels B-H. Note the overall similar distribution and overlap of response types across n = 3 mice. Two US responsive units in mouse FT6963 showed a *reduction* in firing upon the footshock (J, dashed areas).

### Behavior

Behavioral experiments were performed after 3-4 weeks of post-surgical recovery period. Prior to behavior testing, animals were habituated to handling and to the head tethering with the optic patch cords. A classical auditory cued threat memory paradigm was performed in a conditioning chamber of a NIR Video Fear Conditioning Optogenetics Package for Mouse (MED-VFC-OPTO-M, Med Associates Inc., VT, USA). The 3-day protocol consisted of the following steps (see Fig. 3F). On Day 1, the animal was subjected to a habituation session when six tone blocks (CS) were delivered at pseudo-random intervals in a context A. Each tone block consisted of 30 beeps (7 kHz, 80 dB, 100 ms long, repeated at 1 Hz for 30 s). On Day 2 (the same context A), each of the 30s tone blocks was followed by a 1s - long mild electric footshock applied through the metal grid floor of the conditioning chamber (0.6 mA AC). On Day 3, four CS tone blocks were delivered in a new context B. For this, the grid floor was replaced with a smooth white acrylic floor, and a curved wall was installed instead of a square arena used in the context A. The Context A chamber was cleaned before and after each session with 70% ethanol, whereas the Context B chamber was cleaned with a general purpose soap. On some occasions, we tested the retrieval of contextual threat memory on Day 4 by monitoring the animal's freezing behavior in context A for 5 min (Fig. 3F).

**Fig. 3.**
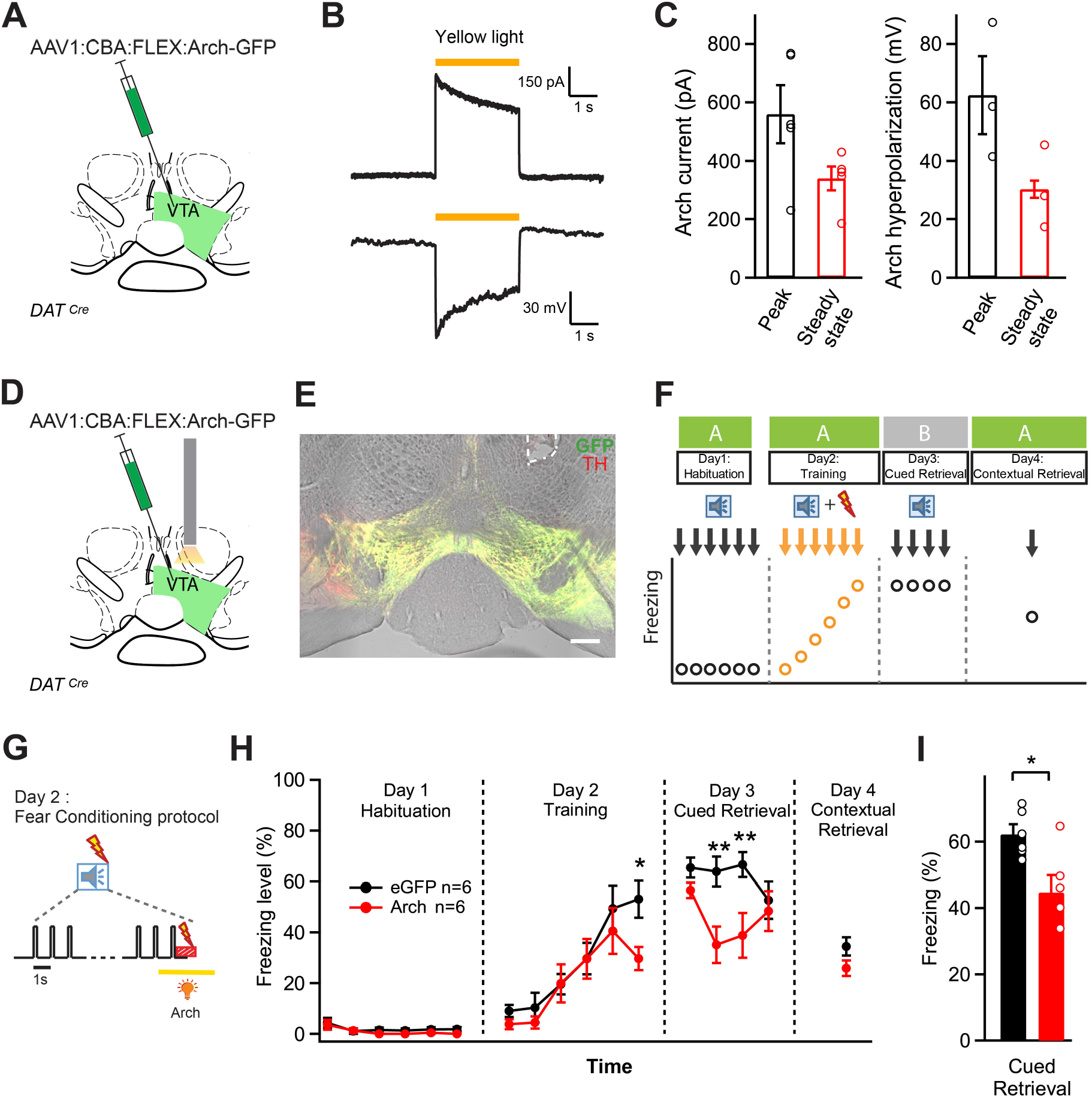
Photoinhibition of VTA dopamine neurons during the footshock decreased the amount of threat memory. (**A**) Schematic of the experimental approach for panels B, C; an AAV1:CBA:FLEX:Arch-eGFP vector was injected unilaterally into the VTA. (**B**) Slice electrophysiology showing current (top; holding potential, −60 mV) and membrane potential hyperpolarization (bottom) evoked by yellow light (◻ = 595 nm). (**C**) Quantification of the peak- and steady-state current (left) and hyperpolarization (right) from n = 5 recordings in n = 3 mice. (**D**) Schematic of the experimental approach for panels E-I. A stereotactic injection of the Cre-dependent Arch-GFP vector into the VTA of DAT^Cre^ mice was followed by unilateral optic fiber implantation above the injection site. (**E**) Post-hoc histological verification of Arch-GFP expression and optic fiber placement (white dashed line) in the VTA of one example mouse. The section were immunolabelled with an TH antibody (red channel), and fluorescence of Arch-GFP construct is shown in green. Scale bar, 200 μm. (**F**) Schematic diagram of the fear learning protocol. See Results and Materials and Methods for details. (**G**) The timing of yellow light application (3s; *λ* = 561 nm) centered over each 1s-footshock stimulus on day 2. (**H**) Average time course of freezing level in Arch - expressing mice (n = 6, red), and in eGFP - expressing control mice (n = 6; black). The stars indicate the significance of the photoinhibition effect as assessed by Bonferroni post-hoc test at the indicated time points, following a two-way repeated measures ANOVA. (**I**) Percentage of freezing during cued threat memory retrieval averaged over four CS presentations on day 3 (n = 6 and n = 6 mice in the test and control group; p = 0.0134, t-test). One data point from a mouse in the control group showed unusually low freezing upon retrieval (26 %) and was removed from the data set.

Freezing was quantified using VideoFreeze® software (Med Associates) from the recorded videos of behaving mice (30 frames/sec). The freezing level (Figs. 2 – 5) was expressed as the percentage of time the mouse spent freezing during the CS presentation (30 s - long tone block).

### Optrode recordings

For recording extracellular spiking activity in the VTA (Fig. 2), we used a 16-channel amplifier (ME16-FAI-μPA; Multi Channel Systems, Germany) and custom-built microdrive mounted optrodes. The general design to build the optrodes followed (Anikeeva et al., 2012); specific manufacturing steps and materials were described in Berret et al. (2019). We used ChR2 expression in DAT^+^ neurons to optogenetically identify recorded units (*DAT*^*Cre*^ × *ChR2-eYFP* mice). In one case a *DAT*^*Cre*^ mouse (FT6963) was injected with AAV8:hsyn:FLEX:ChETA-eYFP; this vector was custom packaged in the lab of Dr. B. Schneider (EPFL).

One day before the behaviour testing, the optrode was re-positioned under ketamine anesthesia (90 mg/g body weight ketamine and 10 mg/g xylazine) to search for opto-tagged units in the VTA by advancing it ventrally from the initial implantation site. The optrode was slowly advanced until light-evoked spikes could be detected at a latency of 2-8 ms (see Fig. 2B-C). Opto-tagging was done by delivering short (2-5 ms, 3-10 mW) light pulses at 2 Hz repetition rate, from a 473 nm DPSS laser (MBL-FN-473-150mW; CNI Lasers). The opto-tagging procedure was repeated after each behaviour session (without advancing the optrode in the later sessions) to collect light-evoked spikes in response to ~2000 light pulses.

Raw data was recorded with a MC_Rack software (MultiChannel Systems) at 40 kHz sampling rate, and was band-pass filtered (0.6-6 kHz) before the analysis. Off-line analysis was performed in IgorPro 7.08 (Wavemetrics, OR, US) as described previously (Berret et al., 2019). Spike waveforms were clustered using a MClust toolbox (Dr. David Redish; University of Minnesota, USA) running KlustaKwik algorithm (Rossant et al., 2016) under MATLAB (MathWorks, USA). The quality of the clusters was assessed by computing the isolation distance (ID) and L-ratio to control for type I and type II clustering errors (Schmitzer-Torbert et al., 2005); only clusters with ID > 24 and L-ratio < 0.5 were retained.

Response types of the units to sensory stimuli were classified according to the following criteria. i) The unit was considered US-responsive if the time-averaged z-score during the 1s - long footshock calculated from the average post-stimulus timing histograms of six US presentations on Day 2 exceeded a value of + 2. ii) For the CS response classification, average z-scores over 100 ms - long tones were calculated from 30 aligned and averaged tone responses. A unit was considered CS-entrained if, *first*, the z-score did not exceed 2 for at least five out of six CS presentations on Day 1 (i.e. the unit was not innately responsive to sound), and, *second*, if out of the last three tone presentations on Day 2, at least two resulted in a summed z-score larger than 4 (i.e. tone response increased on Day 2 as a result of training), *or* if at least two tone presentations on Day 3 resulted in a summed z-score larger than 4 (i.e. the tone response was maintained on Day 3).

### Fiber photometry measurements of dopamine release

For *in-vivo* fiber photometry, an AAV5:hsyn:dLight1.2 vector (Patriarchi et al., 2018) was injected into the left BA of C57Bl6/J mice (250 nl; 4·10^12^ particles / ml; coordinates relative to bregma: AP −1.35 mm; ML −3.3 mm; DV 5.2 mm). Photometry optic fibers were made from a 400 ◻m core / 0.5 NA / 440 ◻m o.d. fiber (FP 400URT; Thorlabs) and a 1.25 mm stainless steel ferrule (SFLC440; Thorlabs).

The optical setup was based on a modified Mightex OASIS tilting microscope (Mightex Systems, Toronto, Canada). Excitation light was generated by two collimated BioLED modules: a 470 nm emitter (BLS-LCS-0470-15-22, Mightex Systems), and a custom fitted 408 nm UV-emitter (LZ4-40UB00-00U8, LED Engin Inc, San Jose, CA, USA). The beams of the two LEDs were combined using a multi-wavelength combiner (LCS series, Mightex Systems) and further coupled via a light guide into a 40 × objective (LUMPlanFL N, Olympus, Tokyo, Japan). The objective formed an image of the end of the 400 μm core optic fiber onto an EM-enhanced CCD camera (C9100-13, Hamamatsu Photonics, Hamamatsu, Japan; operated at 100 frames/sec under the MicroManager software; (Edelstein et al., 2014). A 495 nm dichroic mirror and a 525/50 nm GFP emission filter (Chroma Technology) were used to separate 408 and 470 nm excitation from the fluorescence of the dLight1.2 probe. The LEDs were driven by a two-channel LED driver (BLS-1000-2, Mightex Systems) under control of a Master-8 stimulator (A.M.P.I., Jerusalem, Israel) to generate 7 ms long light pulses at 408 nm, 470 nm plus a dark exposure. The LED power was calibrated to 100 μW at the tip of the implant for both wavelengths. The effective sampling rate after multiplexing the wavelengths was ~33 Hz. The timestamps of LED pulses, CCD exposures, and the TTL signals from the fear conditioning chamber (Med Associates) were recorded by an EPC-10 patch-clamp amplifier (HEKA Elektronik; Germany).

Off-line analysis of photometry signals was done with custom routines in IgorPro (WaveMetrics). Bleaching and other slow drifts in the signal were removed by fitting 3^rd^ - 5^th^ order polynomials to the background-subtracted 470 nm and 408 nm fluorescence traces. The 408 nm fluorescence signal was scaled with the ratio of the standard deviations of the 470 and 408 nm traces, and then subtracted from the 470 nm signal. The resulting corrected 470 nm signal was used to calculate ΔF/F_0_ traces that were subsequently low-pass filtered at 8 Hz for display purpose (Fig. 5).

### Retrograde tracing, immunohistochemistry and histological analysis

For retrograde labeling (Fig. 1G-L), Alexa Fluor-conjugated cholera toxin subunit B (CTB) was used (CTB–Alexa 647; Thermo Fisher Scientific). CTB was unilaterally injected into the BA of a *DAT*^*Cre*^ × *ChR2-eYFP* mouse, at the coordinates from bregma: AP −1.05 mm, ML 3.25 mm, DV −5.18 mm (250 nl of 0.1% dilution in PBS). Seven days after CTB injection, the brains were dissected and processed for histological analysis.

Immunohistochemistry (Fig. 1A-F) and post-hoc histology to confirm the viral targeting and optic fiber placement (Figs. 2–5) were done according to standard procedures after transcardial perfusion of mice with 4% paraformaldehyde. The following primary antibodies were used: chicken anti-GFP (cat. 13970; Abcam; 1:1000 dilution) and rabbit anti-TH (cat. AB152; Millipore; 1:1000 dilution). Secondary fluorescently tagged antibodies were: goat anti-chicken Alexa-488 (cat. A11039) and goat anti-rabbit Alexa-647 (cat. A21244; both from Thermo Fisher Scientific), used at 1:200 dilution. Slices were mounted in Dako fluorescence mounting medium (Dako, Glostrup, Denmark).

Histological sections were imaged at the Bioimaging and Optics Platform (BIOP), EPFL. For cell counting (Fig. 1), the images were taken on an upright LSM 700 confocal microscope (Carl Zeiss) with 20× / 0.8 NA dry and 40× / 1.3 NA oil immersion objectives. Quantification of eGFP and TH colocalization in the VTA (Fig. 1) was performed by using an automated ImageJ routine (provided by Olivier Burri, BIOP, EPFL). For the post-hoc histological analysis (Figs. 2–5), slices were imaged with a stitching fluorescence microscope DM5500 (Leica) using 5× / 0.15 NA or 10× / 0.3 NA objectives, or using a Slide Scanner VS120-L100 (Olympus) with a 10× / 0.4 NA objective.

### Ex-vivo electrophysiology

For slice electrophysiology, 300 μm thick slices containing the VTA were made from 3-4 months old mice previously injected with AAV1:CBA:FLEX:Arch-eGFP, using a Leica VT1000S slicer (Leica Microsystems, Wetzlar, Germany). Whole-cell patch-clamp recordings of VTA neurons were done at room temperature (21 - 23°C) with a K-gluconate based pipette solution, and a standard bicarbonate buffered extracellular. The set-up was equipped with an EPC10/2 patch-clamp amplifier (HEKA Elektronik), and an upright microscope (Axioskop 2, Carl Zeiss, Germany) with a 60× / 0.9 NA water-immersion objective (LUMPlanFl, Olympus, Japan). For activation of Arch (Fig. 3B), yellow light pulses were delivered from a high-power LED (Amber CREE XP-E2 PC, 595 nm; Cree Inc, NC, USA), coupled to the epifluorescence port of the microscope. The LED was controlled with a Cyclops LED driver (Open-Ephys.org; (Newman et al., 2015). The maximal light intensity used in experiments was estimated to be 1.24 mW/mm^2^ at the focal plane. The electrophysiological recordings were analyzed in IgorPro using NeuroMatic plug-in (Rothman and Silver, 2018).

### Experimental design and statistical analysis

No previous sample size calculation was performed. Optogenetic silencing experiments with Arch (Figs. 3H, I; 4C, D; 4G, H) were usually performed in small “cohorts” of n = 3 and n = 3 mice for “control group” and “Arch group”. Usually, repeating such cohorts 2 - 3 times, giving rise to n = 6 - 9 mice in the control group and n = 6 - 9 mice in the Arch group, was found necessary to determine if silencing induced a significant change in the Arch group as compared to the control group (Figs. 3H, I; 4C, D), or not (Fig. 4G, H).

**Fig. 4.**
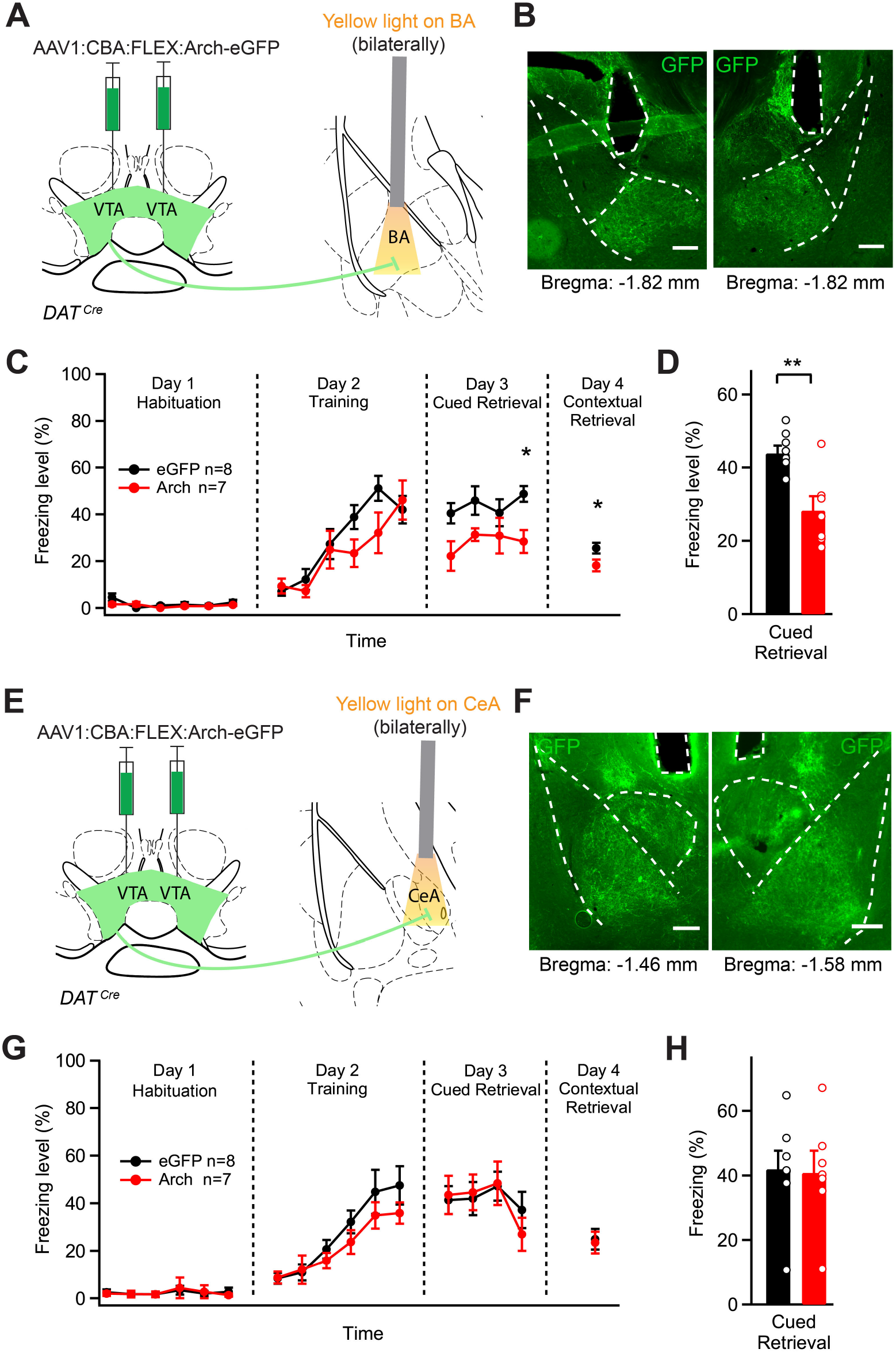
Photoinhibition of the dopaminergic projection from the VTA to the BA during the footshock decreases the amount of cued - and contextual threat memory. (**A**) Schematic of the experimental approach for the panels (B - D). An AAV1:CBA:FLEX:Arch-eGFP vector was injected bilaterally into the VTA of *DAT*^*Cre*^ mice, and optical fibers were implanted above each BA. During the behavioral testing, a 3 s long pulse of yellow light (561 nm) was delivered to each BA starting 1s before the footshock on day 2. (**B**) Post-hoc histological validation of the bilateral optical fiber implantation above the BA (scale bar, 200 μm). (**C**) Average time courses of freezing level in Arch-GFP-expressing (red) and eGFP-expressing control animals (black). The stars indicate the significance of photoinhibition effect assessed by Bonferroni post-hoc test for multiple comparisons at respective time points, following the two-way repeated measures ANOVA. (**D**) Percentage of freezing during cued fear retrieval on day 3, averaged over the four CS presentations (n = 8 and 7 mice in the control and test group respectively; p = 0.0043; t-test). (**E**) Experimental approach for the panels (F - H), in which optical fibers were implanted bilaterally above each CeA. (**F**) Post-hoc histological validation of the bilateral optical fiber implantation above each CeA in one mouse (scale bar, 200 μm). (**G, H**) Time course, and average freezing levels after silencing the VTA dopaminergic fibers over the CeA. There was no significant difference between the Arch and the eGFP (control) group (two-way repeated measures ANOVA in G and t-test in H; p = 0.9). Error bars are mean ± s.e.m.

Replicates often refer to the number of mice or else to the number of cells as indicated in the Results section; in these cases, replicates are biological replicates. Sometimes, technical replicates such as the number of sections analyzed are also indicated in the Results.

Statistical tests were performed using GraphPad Prism 5 (GraphPad Software, CA, USA). The data were expressed as mean ± s.e.m. For the statistical analysis of the optogenetic silencing data (Figs. 3H, 4C, 4G), a repeated measures two-way ANOVA was used, followed by a post-hoc Bonferroni test for multiple comparisons. Additionally, for the analysis of cued retrieval (Figs. 3I, 4D, 4H), the freezing data in response to all n = 4 tone blocks applied on day 3 were pooled, and averaged for each group; in this case, a two-tailed heteroscedastic two-sample Students-t-test was used (referred to as “t-test” in Results). The data in Fig. 5F was analyzed by a one-way repeated measures ANOVA (p < 0.0021; F_(5,25)_=5.19), followed by a post-hoc test for linear trend. The data in Fig. 5G was tested with a one-way repeated measures ANOVA, and with a Dunnett post-hoc test using the first point on Day 1 as a reference. Significance levels are reported in the Results text, and are additionally indicated in the Figures by star symbols according to: p > 0.05, no star; p < 0.05, *; p < 0.01, **; p < 0.001, ***.

**Fig. 5.**
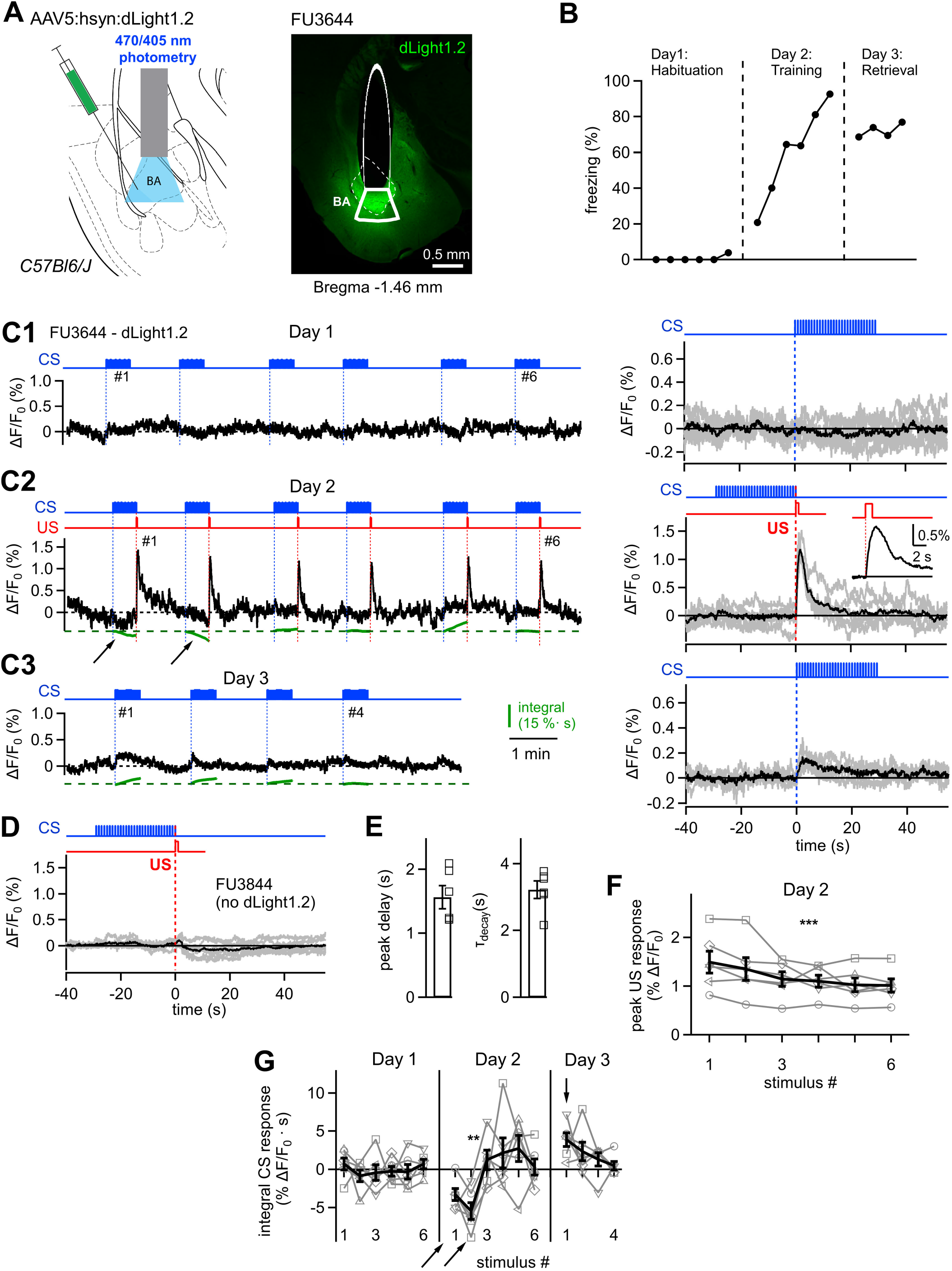
Dopamine release in the BA measured by dLight1.2 during the 3-day fear learning protocol. (**A**) Left, schematic of the fiber photometry experiment with the dopamine sensor dLight1.2. Right, a post-hoc histological image of the dLight1.2 expression (green; enhanced with GFP antibody) in the BA (outlined with dashed line), and of the optic fiber placement above the BA (thin outline shows the fiber body; thick outline shows the estimated illumination area). The image is from the same animal (FU3644) as the example data in panels (B-C). (**B**) Time course of freezing upon tone presentation during the 3 - day fear learning protocol. (**C1-C3**) Left, ◻F/F_0_ traces of dLight1.2 fluorescence recorded in one mouse (FU3644) on habituation day 1 (C1), training day 2 (C2) and threat memory retrieval day 3 (C3). The timing of CS - and US - presentations are shown by blue and red traces, respectively. Robust dopamine concentration transients were observed with every footshock stimulation on day 2 (C2). Integrated ΔF/F_0_ traces for the duration of the CS are shown in green. Right, individual traces of ΔF/F_0_, aligned either to the onset of tone (CS) presentations (C1, C3), or to the onset of the footshock on day 2 (C2). Gray, individual cutout traces; black, average traces. The inset in panel C2 shows the average ◻F/F_0_ trace at higher time resolution. (**D**) Responses to footshocks in a control mouse (FU3844) in which the injection with the AAV5 vector driving the expression of dLight1.2 was omitted. Note absence of ΔF/F_0_ signal (compare to C2, right). (**E**) Average, and individual data points for time to peak (left) and for the decay time constant (right) of dLight1.2 ΔF/F_0_ traces in response to footshocks. N = 6 mice each. (**F**) Average (black) and individual (gray; n = 6 mice) values of peak dopamine concentration transients in response to the n = 6 footshocks on day 2. The responses declined over repeated CS-US pairings on day 2; for statistical testing, see Results. (**G**) Integral ◻F/F_0_ values for CS - responses (integration interval 30s). Gray and black traces are individual and average traces for n = 6 mice. Note the *negative* integral values (arrow) for the first and second CS presentation on day 2 (see also arrows in panel C2), and the positive ΔF/F_0_ value in response to CS stimulation on day 3 (vertical arrow). For statistical testing, see Results. Error bars are mean ± S.E.M.

## Results

### Dopamine neurons of the VTA project to the BA and medial CeA

We first investigated the VTA - amygdala projection anatomically by antero- and retrograde tracing techniques. We used dopamine-transporter Cre mice (DAT^Cre^), and injected an AAV vector that drives the Cre-dependent expression of eGFP into the VTA (AAV1:CAG:FLEX:eGFP; Fig. 1A). Confocal fluorescence images revealed many transduced, eGFP-positive cell bodies within the VTA that were also positive for tyrosine hydroxylase (TH) as revealed by an anti-TH antibody (Fig. 1B). Specifically, we found that 81 ± 9% (1060 / 1259) of all TH - positive (TH^+^) cells in the VTA expressed eGFP (n = 9 sections from n = 3 mice), and that 98 ± 2% (1060/1082) of the eGFP-expressing cells were TH^+^ (n = 9 sections in n = 3 mice; Fig. 1C). Thus, ~ 80% of the VTA neurons were infected with the AAV1 vector, and expression was virtually limited to TH^+^ neurons, demonstrating tight expression control by the Cre-dependent expression vector.

In the amygdala region, we found eGFP - expressing axons in the CeA and in the basolateral nucleus complex (Fig. 1D). In the latter, the highest fiber density was found in the BA, whereas the LA received few eGFP-positive fibers (Fig. 1D). In the CeA, the medial subdivision showed a denser innervation with eGFP - positive fibers as compared to the lateral CeA (Fig. 1D; (Mingote et al., 2015)). Nevertheless, the lateral CeA was densely stained by the TH antibody, consistent with a catecholaminergic innervation of the lateral CeA from a non-VTA source (Asan, 1998; Hasue and Shammah-Lagnado, 2002; Matthews et al., 2016). In the BA and the medial CeA, the expression pattern of eGFP^+^ axons largely overlapped with that of TH (Fig. 1E, F). Besides the BA and medial CeA, TH^+^ fibers, many of which also expressed eGFP, were found in the amygdalostriatal transition zone (AStr) (Fig. 1D). This data shows that DAT-expressing neurons in the VTA send axons to the BA, and to the medial CeA.

We next investigated the location of BA - projecting dopamine neurons in the VTA by using the retrograde tracer Choleratoxin-B (CTB), which was injected into the BA of DAT^Cre^ × ChR2 (Channelrhodopsin-2) mice (CTB-Alexa 647; Fig. 1G). The injections filled a large fraction of the BA but at the same time were limited to the BA (Fig. 1H). Retrogradely labeled neurons were found in the mPFC, the hippocampus and other brain regions known to provide input to the BA (not shown). In the VTA, we similarly observed retrogradely labeled neurons (Fig. 1I, J). We found that BA projectors were homogeneously distributed over the a-p axis (Fig. 1 K, L), and medio-laterally as measured by the distance from the midline (Fig. 1 L). On the other hand, we found more BA projectors in the dorsal half as compared to the ventral half of the VTA (n = 145 of a total of n = 160 neurons were located in the dorsal half; Fig. 1K). Thus, we confirm a preferential dorsal localization of BA projectors within the VTA (Baimel et al., 2017). However, we could not confirm a preferential localization of BA projectors in the more lateral-, and more posterior parts of the VTA; the difference might be caused by the more restricted BA injection volumes in the previous study (Baimel et al., 2017). Taken together, antero- and retrograde labelling experiments reveal a projection from DAT^+^ neurons to the BA and CeA. The projection to the BA is carried by a subpopulation of relatively evenly distributed neurons in the VTA.

### VTA neurons respond to footshock and acquire tone (CS) responses

To investigate the role of VTA dopamine neurons in fear learning, we recorded the activity of VTA dopamine neurons throughout a three-day fear learning paradigm (see Materials and Methods and Fig. 3B for the fear learning protocol). For the *in-vivo* recordings, we used optrodes, which consisted of four tetrode bundles placed around a central optical fiber (Fig. 2A, B) (Anikeeva et al., 2012). Initial attempts to optogenetically identify BA projectors after expression of ChR2 with retrograde viruses in the BA (Herpes Simplex virus) were not fruitful, maybe because of the relatively sparse population of BA projectors in the VTA (Fig. 1). We therefore decided to express ChR2 under the DAT promoter using DAT^Cre^ × ChR2-eYFP mice, which allowed us to identify putative dopamine neurons by optogenetic stimulation (Fig. 2C). The example mouse of the recordings shown in Fig. 2 developed a strong freezing response on the training day, and maintained this level of freezing on the retrieval day (Fig. 2D).

During the training day, when tones were paired with footshock stimulation, a majority of the recorded units showed an increased AP firing in response to the footshocks, and acquired an AP firing response to the tone. The example unit shown in Fig. 2E, F (a putative DAT^+^ dopamine neuron) reliably responded to the footshock (Fig. 2E, right). In addition, this unit developed a short-latency response to the tone beeps when these were followed by a footshock on the training day (Fig. 2E). The tone response was maintained during the retrieval of threat memory one day later (day 3; Fig. 2F). In this mouse, n = 8 units that fulfilled the quality criteria for unit isolation could be followed through at least day 2 and day 3. In n = 4 of these units, a CS response developed during the pairing of the CS with a footshock, whereas other units remained below the criterion for a response (Fig. 2G, pink-versus grey traces, respectively). In the same mouse, n = 6 units responded to footshocks, whereas n = 2 units were non-responsive (Fig. 2H, pink- and gray traces respectively). The AP-firing responses varied in individual units across the six footshocks, but no clear time-dependence could be found (Fig. 2H).

We next analyzed the various response types in the form of Venn diagrams for each mouse (Fig. 2I - K). We made optrode recordings in n = 12 mice, but we could only recover useful single-units that could be followed over at least day 2 and - 3 of the fear learning protocol in n = 3 mice. In these mice, there was a substantial overlap between US-responders and CS-entrained units (Fig. 2I – K; red and blue areas respectively). In all three mice, opto-tagged units (putative DAT^+^ neurons) were a sub-population of US-responding and CS-entrained units (white and yellow areas in Fig. 2I - K), and in each mouse, we found a minority of non-responding units (Fig. 2I - K, gray areas). Furthermore, we observed a *decrease* in AP firing in response to footshocks in n = 1 opto-tagged unit (Fig. 2J, red dashed area), and in n = 1 out of 20 non-identified units (Fig. 2J, blue circle; see Extended Data 1 - Fig. 2 - 1 for the traces).

Taken together, a majority of units recorded in the VTA during the fear learning protocol, amongst them putative DAT^+^ neurons, responded with an increased AP-firing to the footshocks, and acquired a short-latency response to tones when these were reinforced by a footshock. These results suggest that VTA dopamine neurons have a role in auditory-cued fear learning.

### Role of VTA dopamine neurons in auditory threat memory

To investigate whether dopamine neurons in the VTA contribute to the formation of an auditory-cued threat memory, we next employed optogenetic methods. Because we found that many VTA neurons, including dopamine VTA neurons, increase their AP firing in response to footshocks (Fig. 2), we next wished to silence VTA dopamine neurons at the time of the footshock, and observe the effect on fear learning. For this, we employed DAT^Cre^ mice, and expressed the light - activated proton pump Arch-eGFP in a Cre-dependent manner virally in the VTA (using AAV1:CBA:FLEX:Arch-eGFP; Fig. 3A). Initial slice electrophysiology experiments showed that yellow light (561 nm) caused robust outward currents and strong hyperpolarization of Arch - expressing VTA dopamine neurons (Fig. 3B, C; n = 5 recordings from n = 3 mice). We next tested whether VTA dopamine neuron firing during the footshock is needed for threat memory formation. We injected AAV1:CBA:FLEX:Arch-eGFP into the VTA of DAT^Cre^ mice and implanted a single optic fiber over the VTA (Fig. 3D, E). We used a fear learning protocol in which we applied six tone blocks on day 1 without footshock (Fig. 3F; “Day 1”). On day 2, the tone blocks co-terminated with a 1s footshock, and on day 3, retrieval of the auditory-cued threat memory was assessed by applying n = 4 tone blocks alone in a different context (Fig. 3F). To ask whether footshock-driven firing of dopamine neurons contributes to threat memory formation, yellow light was applied for 3s starting 1s before the footshock and ending 1s after the footshock (Fig. 3G). Mice injected with an AAV vector driving the expression of eGFP served as control group.

The average freezing levels during the behavioral tests over all four days of Arch- and eGFP- expressing control mice is shown in Fig. 3H. A two-way repeated measures ANOVA revealed a significant effect of silencing on the freezing levels during the fear learning protocol (p = 0.0117; F_(15, 150)_ = 2.12 for the interaction between the two factors time, and photoinhibition). A post-hoc Bonferroni test showed a significant difference in the freezing level for the sixth tone - footshock pairing on the training day (p < 0.05), and for the second and third tone block on the retrieval day (p < 0.01 for both; Fig. 3H). Pooling the freezing data over the four tone blocks of day 3 indicated a significantly reduced retrieval of cued threat memory (p = 0.013, t-test; Fig. 3I). On the other hand, the contextual threat memory retrieval tested on day 4 was not significantly different between the Arch group as compared to the eGFP group (Fig. 3H, *right*; p = 0.102, t-test). Taken together, these results suggest that footshock - driven activity of VTA dopamine neurons on the training day contributes to the formation of auditory cued threat memory.

### The VTA to BA projection contributes to auditory cued threat memories

The experiments of Fig. 3 suggest that dopamine neurons in the VTA contribute to the formation of auditory - cued threat memories. Nevertheless, there are several possible output projections of VTA dopamine neurons that might contribute to aversive learning, including projections to the mPFC and NAc (Vander Weele et al., 2018; de Jong et al., 2019), as well as the projection to the BA that we characterized anatomically (Fig. 1). To study the role of the dopaminergic VTA to BA projection more specifically, we next silenced the output of the VTA dopamine projections in the BA. This was achieved by bilateral injection of AAV1:CBA:FLEX:Arch-eGFP into the VTA of DAT^Cre^ mice, and implantation of optic fibers bilaterally over each BA (Fig. 4A, B). Mice that expressed eGFP in the VTA but that otherwise underwent identical procedures as the Arch group served as controls. Yellow light (561 nm) was again delivered during a 3s period centered over the footshock stimulation on the training day (Fig. 4C).

In the Arch group, we found a significant decrease of freezing across the fear learning protocol as compared to the eGFP group (Fig. 4C; F_(15, 195)_ = 1.87, p = 0.0238, for interaction between photoinhibition and time; two-way repeated measures ANOVA). Bonferroni post-hoc analysis showed that freezing in response to the fourth tone block on the retrieval day was significantly reduced (p < 0.05). Analysis of the pooled data over all four tone blocks of the retrieval day showed a significant decrease of freezing in the Arch group as compared to the eGFP group (Fig. 4D, p = 0.0043, t-test). Furthermore, contextual threat memory retrieval tested on day 4 was significantly smaller in Arch-eGFP expressing mice as compared to the control group (Fig. 4C, *right*; p = 0.046, t-test). These data show that footshock - driven activity of DAT^+^ axons in the BA originating from the VTA contributes to the formation of auditory cued threat memories, as well as to the formation of contextual threat memories.

Because we found that VTA dopamine neurons also project to the CeA and especially to the medial division of the CeA (Fig. 1; (Mingote et al., 2015), we next tested whether this pathway might contribute to auditory cued threat memory. We expressed Arch-eGFP Cre-dependently bilaterally in the VTA of DAT^Cre^ mice (using AAV1:CBA:FLEX:Arch-eGFP), and implanted optic fibers bilaterally over each CeA (Fig. 4E, F). Yellow light was again applied for 3s during the footshock stimuli. When analyzed over the three subsequent days of the auditory cued fear learning protocol, we found no significant effect of silencing on freezing (Fig. 4G; F_(15, 195)_ = 0.6, p = 0.875 for interaction between photoinhibition and time; two-way repeated measures ANOVA). Similarly, pooling of the freezing data on the retrieval day did not reveal a significant difference between the groups (Fig. 4H, p = 0.90, t-test). Finally, the contextual retrieval of threat memories on day 4 was not significantly different after silencing the dopaminergic VTA axons over the CeA (Fig. 4G, right; p = 0.83, t-test). Taken together, the experiments in Fig. 4 show that the dopaminergic VTA to BA pathway contributes to the formation of auditory cued threat memories and to contextual threat memories, while the VTA to CeA projection seems not necessary for these learning behaviors.

### Footshock stimuli drive dopamine release in the basal amygdala

Previous microdialysis studies have shown that dopamine concentrations rise in the amygdala in response to stressful stimuli including footshocks (Young and Rees, 1998; de Oliveira et al., 2011). Nevertheless, dopamine release has not been measured in the amygdala with high time-resolution at different time points throughout fear learning protocols. We found above that many VTA neurons, including dopaminergic (DAT^+^) neurons respond with an increased AP - firing to footshock stimulation, and acquire responses to tones reinforced by footshocks (Fig. 2). We therefore wished to investigate whether dopamine is released in the BA during footshock stimuli as we would expect, and second, whether dopamine release might develop in response to the learned CS at the time of threat memory retrieval.

To measure dopamine release during the 3-day fear learning protocol in the BA, we used the dopamine sensor dLight1.2 (Patriarchi et al., 2018). We expressed dLight1.2 in the left BA using an AAV5 vector, and measured its fluorescence with an optical fiber implanted over the BA three weeks later during the fear learning protocol. Tone stimulation alone during habituation on day 1 did not notably increase the dopamine concentration in the BA (Fig. 5C1). On day 2, when each tone block was followed by a 1s footshock, we observed robust increases of the dLight1 fluorescence signal, that started precisely with each 1s - footshock stimulus (Fig. 5C2). These dLight1 fluorescent changes were reproducibly observed upon each footshock presentation (Fig. 5C2; Fig. 5F; n = 6 mice), but they were absent in n = 3 control mice in which expression of dLight1.2 was omitted (Fig. 5D). The footshock - evoked dopamine concentration transients reached their peak ~ 1.8 s after the onset of the 1s-long footshock stimulus, and had a decay time constant of 3.2 ± 0.3 s (n = 6 mice; Fig. 5E). The peak amplitude of the dopamine transients decreased with successive footshock stimuli (Fig. 5F; one-way repeated measures ANOVA, [F_(5,25)_=5.19, p = 0.0021], post-hoc test for linear trend, p < 0.0001). The decay of the dLight1.2 peak signal with repeated footshocks might indicate that dopamine responses are stronger with novel, unexpected stimuli (Wise, 2004). A post-hoc histological analysis of fiber position indicated that the amplitude of dLight1.2 transients correlated with the precision of the fiber implant over the BA (Extended Data 2 - Fig. 5 - 1). Taken together, these data show that footshock stimulation during the fear learning protocol induces robust dopamine release in the BA.

We next analyzed the possibility of dopamine release in the BA during the tone presentations. In response to the tone stimulation on day 3, the example mouse of Fig. 5 showed a well - measurable dopamine concentration transient in the BA at the onset of each tone block (Fig. 5C3). Nevertheless, the amplitude of the tone - driven dopamine transients upon threat memory retrieval was smaller than the amplitudes of the footshock - evoked dopamine transients on day 2 (compare Fig. 5C2 with 5C3; p = 0.00071, paired t-test). When we analyzed the integral of the dLight1.2 traces during the 30s tone presentations, a *decrease* in dopamine became apparent in response to the first two CS presentations on day 2 (Fig. 5C2, C3, green traces and arrows; Fig. 5G, arrows). Furthermore, we observed increased dopamine levels in response to the CS presentation on day 3 (Fig. 5G, vertical arrow). A 1-way repeated measures ANOVA showed that the dLight1.2 integral signals in response to the CS underwent a significant time-dependent change (p < 0.0001; F_(15,75)_=4.62; n = 6 mice). A post-hoc Dunnett test showed that the integral dLight1.2 signal in response to the second CS presentation on day 2 was significantly smaller than the response to the first CS presentation (Fig. 5G, p < 0.01). On the other hand, the increased dLight1.2 response to the first CS presentation on day 3 was not significant (p > 0.05; post-hoc Dunnett test; see Fig. 5G, vertical arrow). Taken together, dLight1.2 photometry shows that footshock stimuli cause robust dopamine release in the BA during the training day of the fear learning protocol, whereas dopamine responses to the learned CS are smaller, and more variable between mice.

## Discussion

Using *in-vivo* optogenetic methods and optrode recordings, as well as anatomical circuit tracing and *in-vivo* imaging of dopamine release, we have studied the role of the VTA to BA dopamine pathway in the formation of an auditory cued threat memory. Silencing the VTA - BA pathway during footshock presentation led to a reduced expression of threat memory one day later, suggesting that footshock-driven activity of VTA axons in the BA facilitates the formation of an auditory cued threat memory. *In-vivo* optrode recordings of single units in the VTA showed that most recorded units, including putative dopamine neurons, responded to the footshock with increased AP firing rates, and acquire CS responsiveness. Furthermore, dLight1 imaging showed that footshocks, and to a smaller degree the learned CS, cause dopamine release in the BA. Thus, we conclude that VTA dopamine neurons contribute to signal salient somatosensory events to the BA, where the resulting dopamine release facilitates the formation of an auditory cued threat memory.

A role of dopamine in aversive learning has first been hypothesized based on pharmacological experiments with D1- and D2-receptor blockers and microdialysis - based dopamine measurements in the amygdala (Lamont and Kokkinidis, 1998; Young and Rees, 1998; Guarraci et al., 1999; Nader and LeDoux, 1999). Furthermore, fear-potentiated startle was reduced in the dopamine-deficient mouse model and after conditional genetic deletion of NMDA-receptors in TH^+^ neurons (Fadok et al., 2009; Zweifel et al., 2011). On the other hand, the generally accepted role of VTA dopamine neurons in reward processing (Schultz, 1998; Wise, 2004), and recordings suggesting that VTA dopamine neurons might be uniformly *inhibited* by aversive stimuli (Ungless et al., 2004; Tan et al., 2012), might have hampered progress in understanding the role of dopamine in aversively motivated learning. More recent circuit tracing analyses with viral approaches, as well as retrograde labeling and electrophysiology after behavioral tests showed that the VTA is composed of neurons with diverse projection areas belonging to different subsystems (Lammel et al., 2011; Lammel et al., 2012; Beier et al., 2015). These dopaminergic subsystems in the VTA can be involved in both reward - based learning (for most NAc projectors) or in aversive learning (mPFC projectors). Nevertheless, the VTA to amygdala pathway had received little attention in recent studies.

We made *in-vivo* optrode recordings in the VTA throughout the 3-day fear learning protocol. Although the yield of the recordings was low, the results showed that in behaving mice, most recorded units, including putative dopamine neurons identified optogenetically under the DAT promoter, were excited by the footshock (Fig. 2). We only found few units that were inhibited by the footshock (~ 10%), even within the sample of putative DAT^+^ neurons (Fig. 2I-K). Furthermore, we found that a population of units which largely overlapped with the US - responsive population developed a response to tones when these when re-inforced by the footshock (Fig. 2). It was shown, largely based on AP - waveform criteria, that footshocks in anesthetized mice excite putative VTA GABA neurons, and mostly inhibit putative VTA dopamine neurons (Tan et al., 2012). It remains possible that a fraction of the non-identified units in our sample represented GABA neurons. On the other hand, we showed with optogenetic tools that inhibition of VTA dopamine neurons during the footshock (using DAT^+^ Cre mice), or inhibition of VTA dopamine axons over the BA, significantly reduced threat memory formation (Figs 3, 4). These results, and our finding of robust, footshock-triggered dopamine release in the BA (Fig. 5), strongly suggest that footshocks activate dopamine neurons in the VTA, and that the resulting release of dopamine in the BA facilitates threat memory formation.

Several recent studies have provided evidence for roles of distinct dopamine neuron populations, both within and outside the VTA, in aversively motivated learning. One study found that VTA dopaminergic axons that project to a ventro-medial area of the NAc shell were excited by footshocks (de Jong et al., 2019). Interestingly, these axons acquired a tone response when the latter was re-inforced by footshocks, similar to what we found here for a large fraction of recorded VTA units (Fig. 2). De Jong et al. (2019) also showed that projections to the lateral shell of the NAc – these dopamine axons were involved in reward processing - were *inhibited* by footshocks. Yet another population of VTA dopamine neurons, the mPFC projectors, are involved in aversive valence processing (Lammel et al., 2011; Vander Weele et al., 2018), and a dopamine projection from the VTA to the NAc and to the CeA is involved in fear memory and fear discrimination (Jo et al., 2018). Thus, together with the VTA dopamine neurons projecting to the BA that we have studied here, there are different populations of VTA dopamine projection neurons involved in aversive learning. Furthermore, dopamine neurons located in the dorsal raphe that project to the CeA, and dopamine neurons of the lateral portion of the substantia nigra that project to the posterior striatum, were recently shown to be involved in aversive learning (Groessl et al., 2018); (Menegas et al., 2018). The molecular logics of these dopamine neuron populations with distinct projections patterns has recently been addressed by using intersectional genetic approaches (Poulin et al., 2018). Together, these functional - and genetic circuit studies corroborate the anatomical notion that distinct populations of midbrain dopamine neurons have largely separate projection targets (Swanson, 1982; Menegas et al., 2015). The various dopamine projection pathways that are involved in aversive learning (see discussion above) might be regarded as parallel streams of neuromodulatory input to largely separate projection domains in the telencephalon. The functional advantage for such parallel streams of dopamine signaling should be further elaborated in future work.

We have found a contribution of the VTA - BA dopamine projection to the formation of auditory cued threat memories (Figs 3, 4), and our finding that dopamine is released precisely at the start of the footshock (Fig. 5) supports the notion that dopamine can contribute to a “teaching” signal for plasticity in the BA (Herry and Johansen, 2014). On the other hand, the LA, which receives many sensory inputs from cortical and thalamic areas (LeDoux et al., 1990; Nabavi et al., 2014; Lucas et al., 2016), was only sparsely innervated by dopamine fibers from the VTA (Fig. 1). It therefore remains unclear whether dopamine plays a role for associative plasticity in both the LA and the BA, or whether the role of dopamine is limited to the BA. What might be the mechanisms by which dopamine acts in the BA to promote threat memory formation? Dopamine acts on metabotropic D1- and D2-like receptors (Neve et al., 2004; Tritsch and Sabatini, 2012), and activation of both D1- and D2- receptor subtypes in the amygdala (Lamont and Kokkinidis, 1998; Guarraci et al., 2000) is expected to modulate various signaling aspects. These include an up-regulation of intrinsic neuronal excitability via D1-receptors (Kroner et al., 2005; Tang, 2018), a D2 - receptor mediated decrease in transmission at both excitatory- and inhibitory synapses (Rosenkranz and Grace, 1999); (Chu et al., 2012), and an action of D1- receptors on intracellular signaling pathways involved in long-term plasticity of excitatory synapses (Li et al., 2011). Thus dopamine likely facilitates the induction of plasticity at excitatory synapses in an “instructive” manner involving the activation of D1-receptors, or more indirectly in a “permissive” manner via D2-receptor mediated inhibition of GABA release onto BA principal neurons (Bissiere et al., 2003; Chu et al., 2012).

In summary, our study shows that many VTA neurons, and amongst them dopamine neurons, are excited during footshock stimulation and aversive learning. The resulting dopamine release in the BA facilitates the formation of an auditory - cued threat memory. It will be interesting to investigate which specific excitatory, or inhibitory pathways in the BA are modulated by dopamine to facilitate plasticity, and ultimately the formation of a threat memory. Understanding how dopamine acts as a teaching signal in aversively motivated learning will allow us to gain further insights into the mechanisms of maladaptive plasticities that underlie anxiety and post-traumatic stress disorder (Yeh et al., 2018).

## Acknowledgments

We thank Mrs. Heather Murray, Mrs. Tess Baticle and Mrs. Jessica Dupasquier for expert technical assistance, and Dr. Bernard Schneider (EPFL) for help with AAV vector packaging. Image acquisition was done at the Bioimaging & Optics Platform of EPFL (BIOP). This work was supported by a grant form the Swiss National Science Foundation (SNSF; 31003A_176332 / 1 to R.S.), by the SNSF National Competence Center for research Synapsy - The synaptic bases of mental disease (project #28, to R.S.), and by an EMBO fellowship (ALTF 224-2015; to M.K.).

---

**Extended Data 1 - Fig. 2 - 1.**
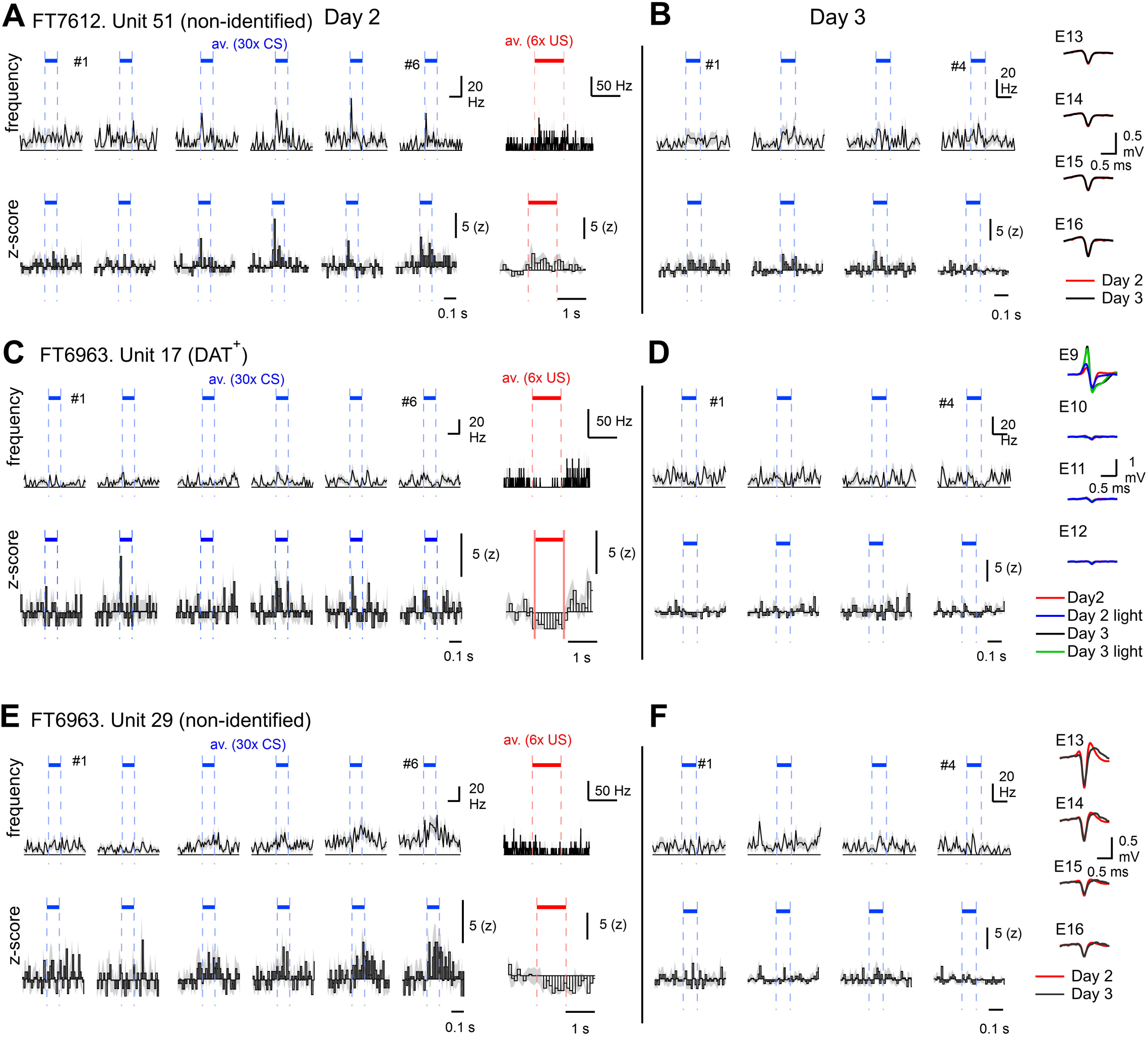
Examples of spiking activity of opto-tagged (DAT^+^) and unidentified single units. The data is presented in a format identical to Fig. 2E, F; stimulus-aligned AP-firing frequency (top row) and z-score (bottom) are shown for the training day 2 (CS presentations in blue, US in red) and for the retrieval day 3 (for CS only). Average AP - waveforms for the four electrodes are shown on the right. (**A-B**) Spiking activity of an unidentified unit from the same example animal as shown in Fig. 2B-H (mouse FT7612). The unit was classified as non-responsive to US, but as CS-entrained (note an increase in CS-evoked spiking over repeated CS - US pairings on day 2). (**C-D**) A DAT^+^ unit from another mouse (FT6963) which showed no CS entrainment, and reduced AP – activity in response to the footshock (C). This unit is highlighted by the red dashed area in Fig. 2J. (**E-F**) Example for a non-identified unit measured in mouse FT6963. This unit also showed reduced AP – firing activity in response to the footshock (see E), and showed CS-entrainment during day 2. This unit is highlighted by the blue dashed circle in Fig. 2J.

**Extended Data 2 - Fig. 5 - 1.**
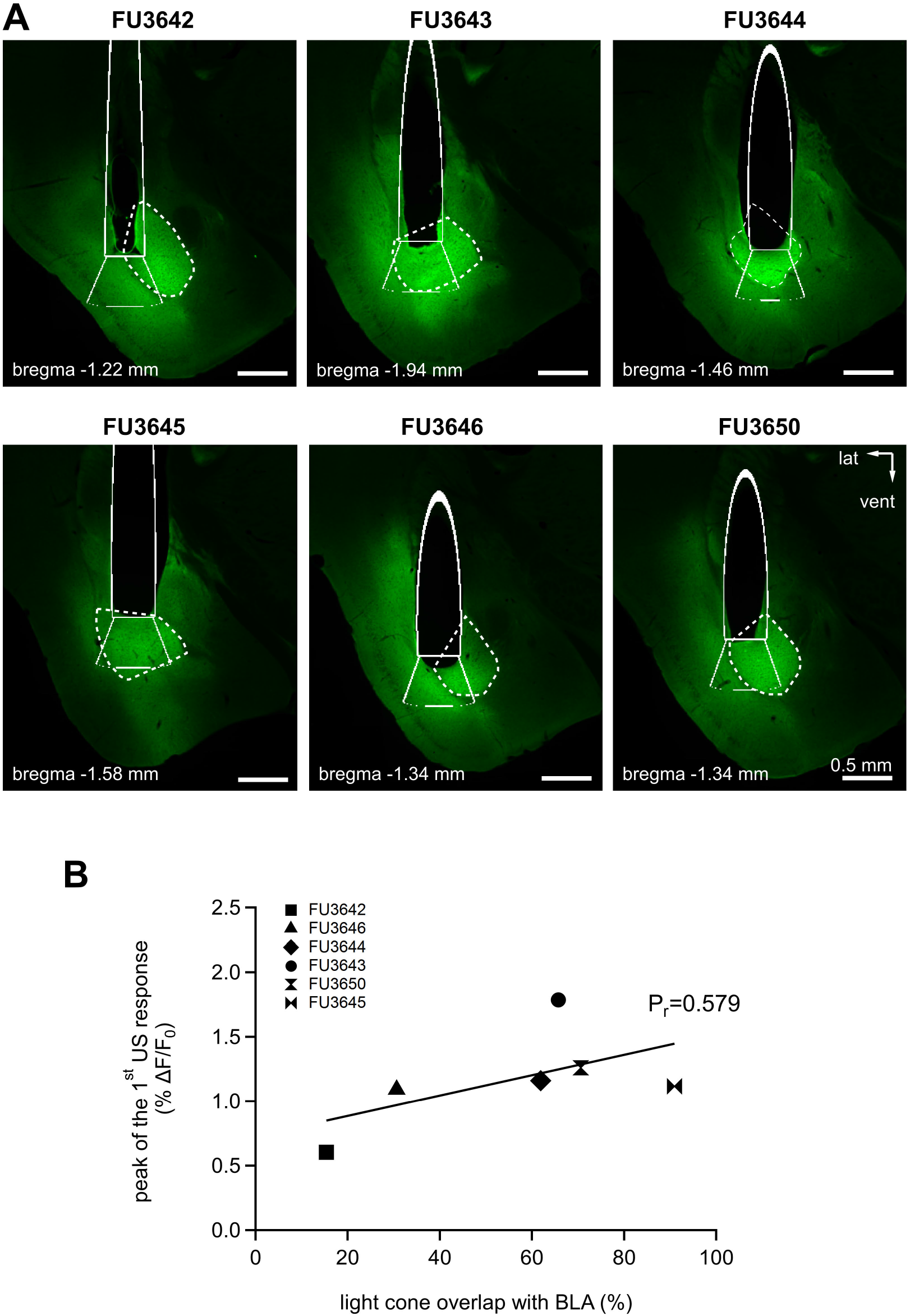
Post-hoc histological verification of optic fiber placement for dLight1.2 measurements of dopamine release in the BA. (**A**) Immunofluorescent images of coronal sections through the left BA of n = 6 mice showing expression of the dLight1.2 probe (green; detected with an anti-GFP antibody). The outline of BA is shown with a dashed line. The solid line represents the reconstructed cross-section through the photometry optic fiber, with an emanating light cone at the ventral end of the fiber (a 500 μm effective illumination radius was assumed; see Aravanis et al., 2007). (**B**) The peak ΔF/F_0_ values of the first US responses of dLight1.2 probe on the training day 2 (see Fig. 5C2) were plotted versus the percent area of the light cone profile that overlapped with the outline of the BA nucleus. The overlap area was summed up over serial brain sections covering the full extent of the light cone in 3D. Note that in mice with lower overlap (e.g. FU3642, FU3646; see A) the US-triggered ΔF/F_0_ responses were lower than in the mice with larger overlap (Pearson’s correlation coefficient of 0.579).

